# A reverse Turing-test for predicting social deficits in people with Autism

**DOI:** 10.1101/414540

**Authors:** Baudouin Forgeot d’Arc, Marie Devaine, Jean Daunizeau

**Affiliations:** Département de Psychiatrie, Université de Montréal, Canada; Centre Intégré Universitaire de Santé et Services Sociaux de Nord-de-l’Île-de-Montréal; Université Pierre et Marie Curie, Paris, France; Institut du Cerveau et de la Moelle épinière, Paris, France; INSERM UMR S975

**Keywords:** social cognition, mentalizing, theory of mind, flexibility, computational psychiatry

## Abstract

Social symptoms of autism spectrum disorder (ASD) are typically viewed as consequences of an impaired Theory of Mind, i.e. the ability to understand others’ covert mental states. Here, we test the assumption that such “mind blindness” may be due to the inability to exploit contextual knowledge about, e.g., the stakes of social interactions, to make sense of otherwise ambiguous cues (e.g., idiosyncratic responses to social competition). In this view, social cognition in ASD may simply reduce to non-social cognition, i.e. cognition that is not informed by the social context. We compared 24 adult participants with ASD to 24 neurotypic participants in a repeated dyadic competitive game against artificial agents with calibrated mentalizing sophistication. Critically, participants were framed to believe that they were competing against humans (social framing) or not (non-social framing), hence the “reverse Turing test”. In contrast to control participants, the strategy of people with ASD is insensitive to the game’s framing, i.e. they do not constrain their understanding of others’ behaviour with the contextual knowledge about the game (cf. competitive social framing). They also outperform controls when playing against simple agents, but are outperformed by them against recursive algorithms framed as human opponents. Moreover, computational analyses of trial-by-trial choice sequences in the game show that individuals with ASD rely on a distinctive cognitive strategy with subnormal flexibility and mentalizing sophistication. These computational phenotypes yield 79% diagnosis classification accuracy and explain 62% of the severity of social symptoms in people with ASD.

## Introduction

The diagnosis of autism spectrum disorders or ASD is based on alterations in two cognitive domains [1]: reciprocal social interaction (social deficits) and flexibility of behaviour (non-social deficits). Although one of the most heritable neurodevelopmental conditions [2,3], there is a remarkably small overlap between the genes that are associated with the distinct ASD-like behavioural traits in the general population [4]. This may explain ASD’s high clinical heterogeneity [5], and eventually challenge the relevance of research agendas aimed at identifying a unique aetiology for both social and non-social deficits in ASD [6]. Somewhat paradoxically, this also highlights the importance of forging social and non-social neurocognitive theories, which can bridge the gap between biological and clinical observations in ASD [7,8]. This work is a step forward in this direction.

One of the most influential theories about ASD social deficits asserts that these are eventually due to an underlying impairment in Theory of Mind or ToM [9,10], i.e. the ability to understand others’ covert mental states. This has been repeatedly evidenced in children using tests of ToM, e.g., false belief understanding [11–13], sarcasm/irony detection [14,15] or moral evaluation [16,17]. However, these tests yield quite unreliable results in older individuals. For example, high-functioning ASD adolescents and adults successfully pass false-belief [18] or facial emotion recognition [19] tests. Further refinements of the “mind blindness” theory of ASD thus suggest that although adults with ASD may succeed in simple mindreading tasks when explicitly instructed to mentalize, they lack some form of implicit and spontaneous ToM [20–22]. Alternatively, one may argue that social deficits in adults with ASD may only become apparent when the task mirrors the demands of ecological social exchanges, which critically rely on highly contextual and interactive signalling [8,23]. This is because social deficits in ASD may be less about inaccurate processing of “socially-salient” stimuli (e.g., facial expressions, speech prosody, etc…), than about the inability to exploit contextual knowledge about, e.g., the stakes of social interactions, to make sense of otherwise ambiguous cues (e.g., idiosyncratic responses to social competition). In this view, social cognition in ASD may simply reduce to non-social cognition, i.e. cognition that is not informed or constrained by the social context.

Recent advances in artificial social cognition [24–26] now enable us to mimic the way human players adapt to others in the context of simple repeated dyadic games. Rather than asking whether this type of algorithm passes the Turing test [27], we ask whether believing it is human or not changes the way people interact with it. The nature of such “reverse Turing test” will be clearer below (see Figure A0 in the Supplementary Text S1). We start with the premise that social interactions induce a specific evolutionary challenge, namely: forecasting others’ overt behaviour from learned associations with predictive cues, including past behaviour [28,29]. Critical here is the notion that people may engage with others equipped with a cognitive repertoire composed of *many learning strategies,* each of which may be tied to distinct representations and/or policies. Arguably, somewhere at the end of the spectrum lie ToM-related learning strategies that derive from adopting the intentional stance [26,30– 32], whose sophistication increases with the depth of recursive beliefs (as in “I believe that you believe that I believe…”). Nevertheless, learning in social contexts can take less sophisticated forms, ranging from simple heuristics, to trial-and-error learning, to cognitive shortcuts of ToM that simply care about others’ overt reaction to one’s own actions [33]. The ability to compliantly draw learning strategies from one’s cognitive repertoire is what we term flexibility. Importantly, mathematical modelling can be used to turn a given learning strategy into a learning rule (i.e. the precise way in which agents adapt to the history of past actions and outcomes). In appropriate experimental contexts (e.g., dyadic games), this endows learning strategies with a specific behavioural signature that can be disclosed from quantitative analyses of trial-by-trial choice sequences [26]. One can then measure and compare the computational properties of people’s learning repertoire, in particular: its ToM-sophistication and its flexibility. In what follows, were refer to these as “computational phenotypes” of social cognition. We then ask whether people with and without ASD differ with respect to these computational phenotypes, which we infer from observed trial-by-trial choices in dyadic interactive games against artificial players. Critically, participants are not told about the algorithmic nature of their opponents. Rather, we have them believe either that they are competing against each other (social framing) or that they are gambling like in a casino (non-social framing). We focus on peoples’ ability to alter their behavioural strategy as a function of whether or not they think they are competing against someone, hence the “reverse Turing test”. We predict that, in contrast to control participants, adults with ASD would not be able to constrain their understanding of others’ behaviour with the contextual knowledge about the game (cf. competitive social framing), hence failing our “reverse Turing test”.

## Results

We asked 24 adult participants with ASD and 24 control participants to play repeated games against artificial “mentalizing” opponents, which differ in their ToM sophistication (hereafter: *k*-*ToM* agents, see below). In total, each participant played 4×2×2=16 games (4 opponent types, 2 framing conditions, 2 repetitions), where each game consisted in 60 successive trials. To succeed, subjects had to anticipate and predict the behaviour of their opponent, who hid himself in one out of two possible locations at each trial (see Figure 1 below).

**Figure 1:**
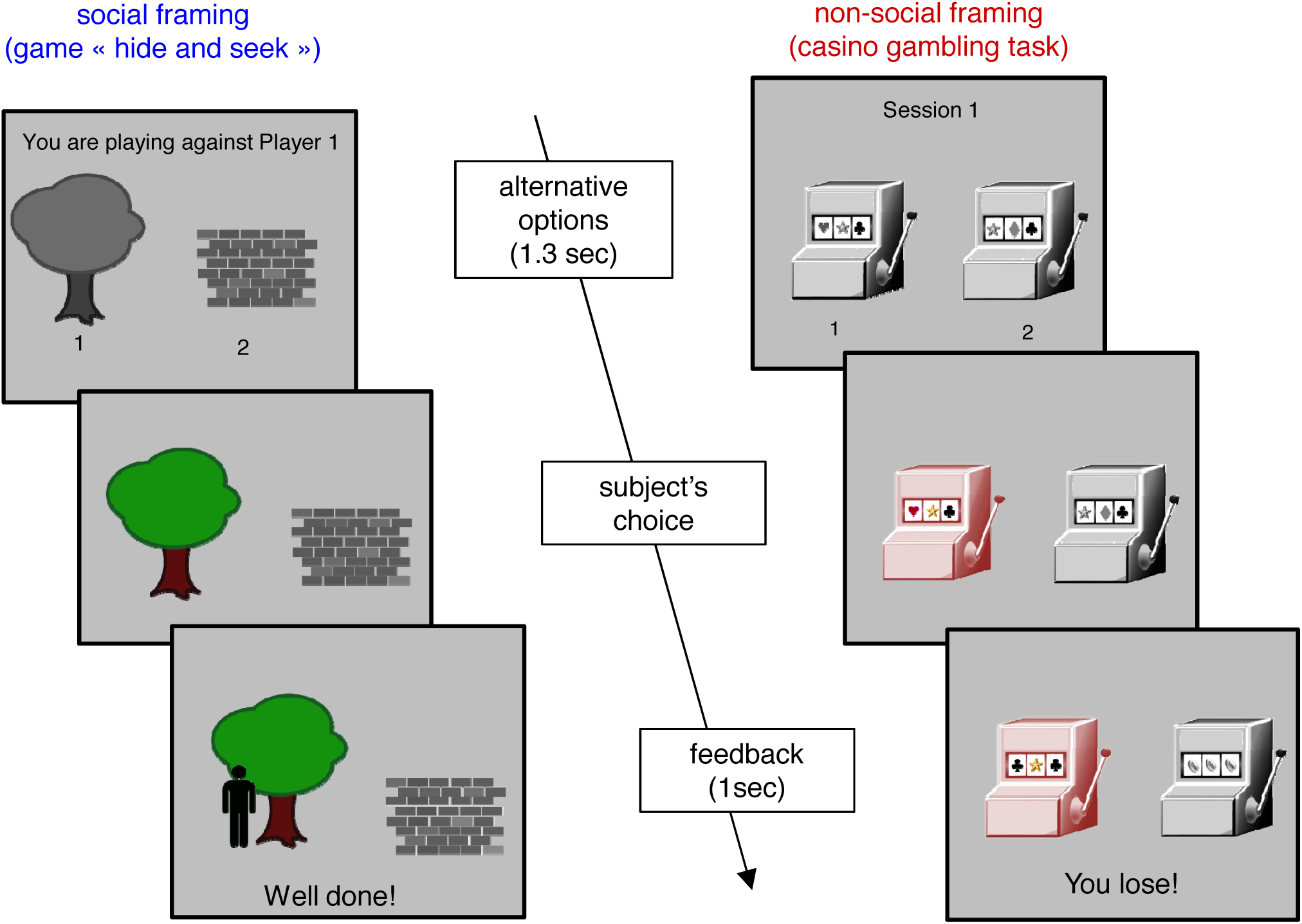
Experimental protocol. Left: social framing (“hide-and-seek” game). Right: non-social framing (Casino game). At each trial, participants have 1300 msec to pick one of the two options (social framing: wall or tree, non-social framing: left or right slot machine). Feedback is displayed for 1 sec; and includes the trial outcome (win or loss) and the actual winning option (social framing: character picture, non-social framing: three identical items).

Opponents either followed a predetermined pseudo-random sequence with a 65% bias for one hand (RB), or were designed to deceive the participants from learned anticipations of their behaviour (*0-ToM, 1-ToM and 2-ToM*). The difference between *k-ToM* opponents lies in how they learn from the past history of participants’ actions, where *k* refers to their calibrated ToM sophistication. In brief, *0-ToM* does not try to interpret the participants’ action sequence in terms of a strategic attempt to win. Rather, it simply assumes that abrupt changes in the participants’ behaviour are a priori unlikely. It thus tracks the evolving frequency of participants’ actions, and chooses to hide the reward where it predicts the opponent will not seek. It is an extension of “fictitious play” learning [34], which can exploit participants’ tendency to repeat their recent actions. In contrast, *1-ToM* is equipped with (limited) artificial mentalizing, i.e. it attributes simple beliefs and desires to participants. More precisely, it assumes that participants’ actions originate from the strategic response of a *0-ToM* agent that attempts to predict its own actions. Note that the computational sophistication of artificial mentalizing is not trivial, since *1-ToM* has to explicitly represent and update its (recursive) belief about its opponents’ beliefs. Practically speaking, *1-ToM* learning essentially consists in an on-line estimation of *0-ToM*’s parameters (e.g., learning rate and behavioural temperature) given the past history of both players’ actions. This makes *1-ToM* a so-called “meta-Bayesian” agent [26,35] that can outwit strategic opponents that do not mentalize when competing in the game (such as 0-ToM). Although 1-ToM is mentalizing, it is not capable of dealing with other mentalizing agents. This is the critical difference between *1-ToM* and *2-ToM*. At this point, suffices to say that *2-ToM* is an artificial mentalizing agent that can learn to predict how other mentalizing agents (such as *1-ToM*) will behave.

Critically, participants were not cued about opponent conditions. This implies that they had to adapt their behaviour according to their understanding of the history of past actions and outcomes. In addition, except in the control (RB) condition, there is no possibility to learn the correct answer from simple reinforcement. This is because *k-ToM* artificial learners exhibit no systematic bias in their response. Further details regarding the experimental protocol as well as k-ToM artificial agents can be found in the Methods section below.

Figure 2 below summarizes the performance results, in terms of the net rate of correct answers in each of 4×2 conditions, for both (control and ASD) groups.

**Figure 2:**
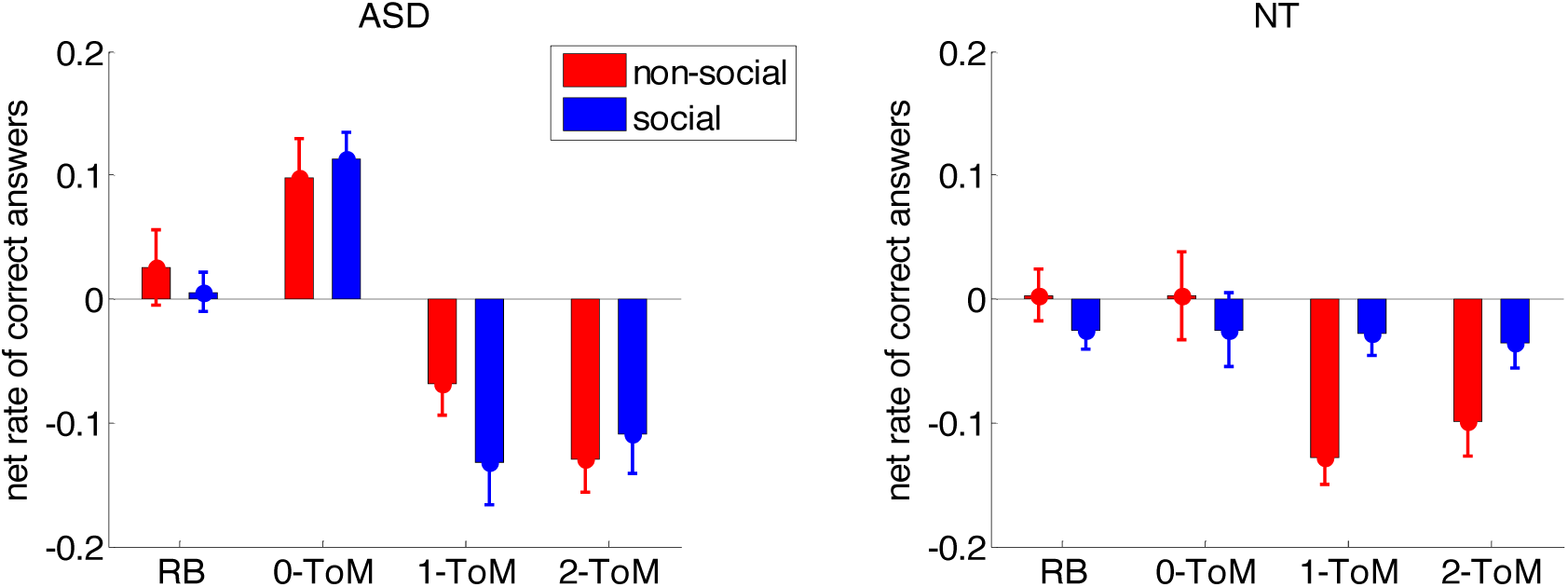
Behavioural performance results. Group average net rate of correct answers (y-axis) against the four opponent types (x-axis) for both framing conditions (blue: social, red: non-social) in both ASD (left) and control (right) participants. Note: The net rate of correct answers is defined as (n+-n-)/(n++n-), where n+ and n-be the number of correct and incorrect responses, respectively. In this and all subsequent figures, error bars depict the standard error around the mean.

One can see that the performance patterns are markedly different between NT and ASD participants. To begin with, the performance of NT participants qualitatively reproduces previous experiments with healthy human adults [26]. In brief, in the non-social framing condition, NT participants eventually lose against artificial mentalizing agents (*1-ToM and 2-ToM*) whereas they maintain their earnings in the social framing condition. The ASD group however, seems to show no effect of the framing manipulation, i.e. their performance pattern across opponents is the same, irrespective of whether they believe that they are competing against other people or not. Interestingly, they seem to lose against artificial mentalizing agents (as NT controls in the non-social framing condition), but they outperform NT controls against non-mentalizing learning agents (*0-ToM*). We performed a pooled variance ANOVA to assess the statistical significance of these observations. We found a significant three-way interaction between group (ASD vs NT), opponent and framing (F[3,690]=3.6, p=0.014, R^2^=1.5%), a significant interaction between group and opponent (F[3,690]=9.5, p<10^-4^, R^2^=4.0%) and a main effect of opponent (F[3,690]=33.7, p<10^-4^, R^2^=12.8%). We then looked more closely at the three-way interaction using post-hoc tests. In the NT group, there was a main effect of opponent (F=4.5, p=0.004), no main effect of framing (F=2.6, p=0.11) but a significant interaction opponent x framing (F=3.7, p=0.011). In the ASD group, there was a main effect of opponent (F=38.7, p<10^-4^) but no main effect of framing (F=0.5, p=0.46) nor interaction (F=1.3, p=0.27). In other terms, only NT participants show the opponent x framing interaction. This is due to the fact that NT participants perform better in the social than in the non-social framing only against artificial mentalizing agents (p<10^-4^). Now focusing on performances against artificial mentalizing agents, there was a significant interaction between group and framing (p=0.001). This is because against *1-ToM* and *2-ToM*, NT participants perform significantly better than ASD people against artificial mentalizing agents in the social framing (p<10^-4^) but not in the non-social framing (p=0.65). Besides, ASD participants perform significantly better than NT participants against *0-ToM* (p<10^-4^), and this effect does not depend upon the game’s framing (p=0.46).

At this point, we asked whether we could classify ASD and NT participants based upon their performance patterns in the task. Averaging performances over repetitions yielded a feature space of 8 dimensions (4 opponent types, 2 framings), which was then fed to a classifier based upon logistic regression [36]. Test classification accuracy was evaluated using a simple leave-one-out cross-validation scheme. The classifier achieved 73% of correct out-of-sample classifications, which is statistically better than chance (p=0.001). This will serve as a reference point for evaluating the added-value of computational phenotypes.

One of the main differences between NT and ASD participants is thus that the latter seem to be insensitive to the framing manipulation. This interpretation, however, neglects the possibility that distinct leaning strategies may eventually yield similar performances in the game. In other terms, performance measures are potentially blind to learning strategies, which can only be inferred from analyses of trial-by-trial action sequences in the game. We thus considered a set of eight distinct learning models that constitute peoples’ potential learning repertoire. Each of these learning models provides a probabilistic prediction of observed peoples’ trial-by-trial choice sequences. We then performed a subject-specific bayesian model comparison of these models, and evaluated both the flexibility -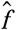- and the ToM-sophistication -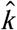- of peoples’ learning repertoires. In what follows, we refer to these as peoples’ “computational phenotypes”. We refer the interested reader to the Methods section.

We first asked whether control and ASD participants would show differences in their repertoire’s ToM-sophistication. Figure 3 below shows the repertoire’s ToM-sophistication 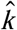averaged across repetitions, across opponent conditions and across participants, for each group and for both framing conditions.

**Figure 3:**
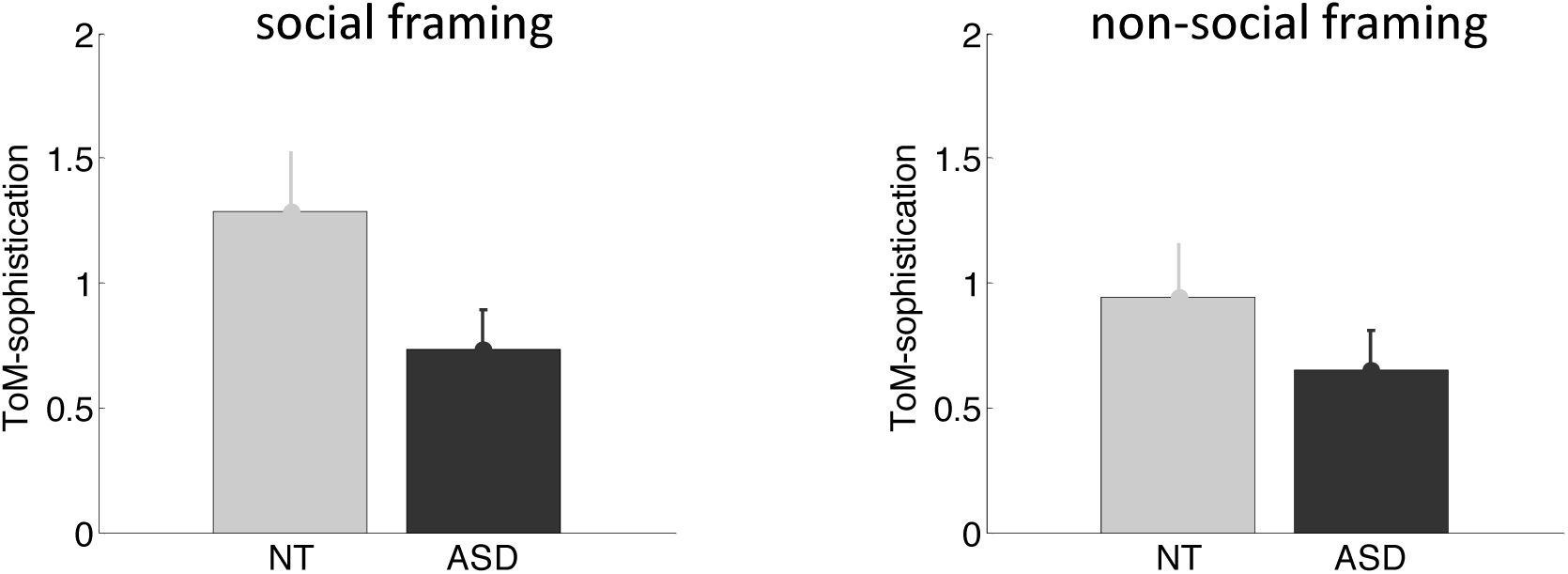
Model-based analysis of trial-by-trial choice sequences: ToM sophistication scores. ToM sophistication scores are shown as a function of framing conditions (left: social, right: non-social) for both control (gray) and ASD participants (back).

A simple ANOVA shows no evidence for an interaction between group and framing (F[1,46]=0.6, p=0.42, R^2^=1.4%), no main effect of framing (F[1,46]=1.8, p=0.18, R^2^=3.8%), but a significant group effect (t[46]=1.9, p=0.03, R^2^=7.3%). Post-hoc tests show that this group difference is mostly driven by the social framing condition: whereas there is no significant difference between the groups in the non-social condition (t[46]=1.1, p=0.13, R^2^=2.7%), there is a strong group difference in the social framing condition (t[46]=1.9, p=0.03, R^2^=7.5%). In other words, only in the social framing do control participants exhibit higher ToM-sophistication than ASD participants.

We then investigated whether control and ASD participants show differences in their repertoire’s flexibility. Figure 4 below shows the repertoire’s flexibility 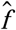, both across framings and across repetitions.

**Figure 4:**
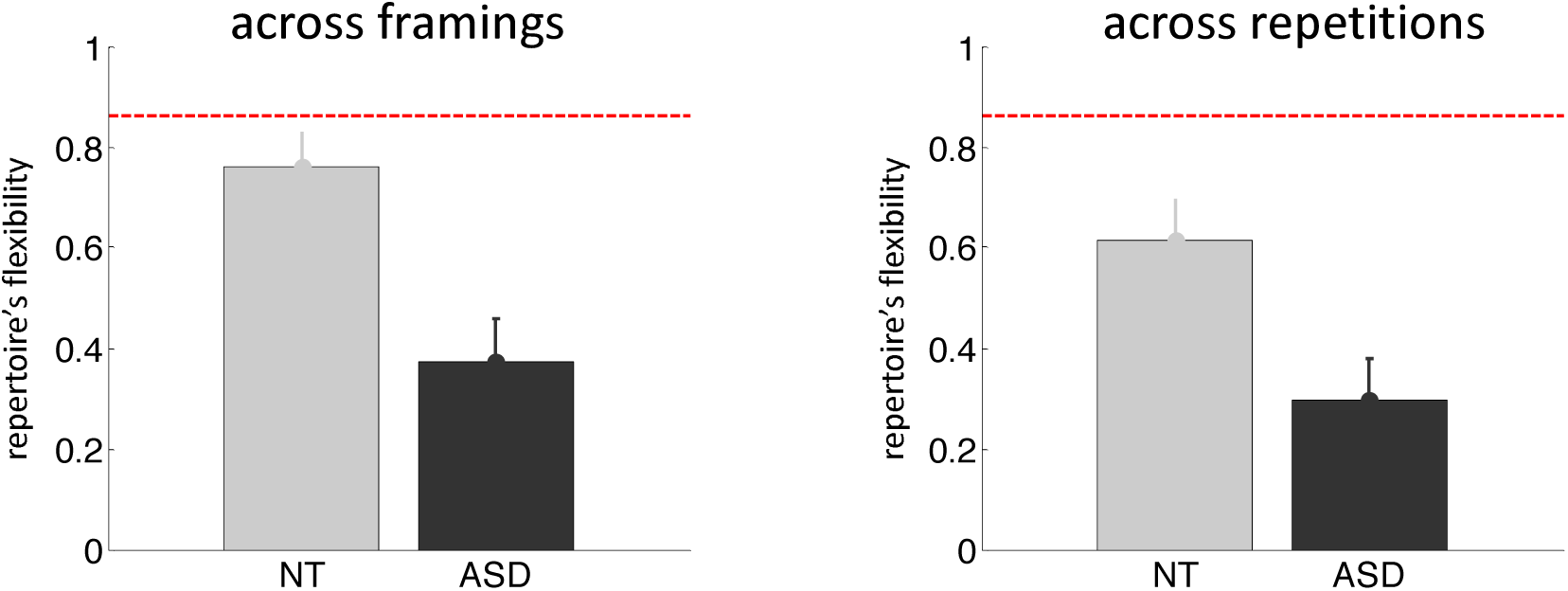
Model-based analysis of trial-by-trial choice sequences: repertoire’s flexibility. The repertoire’s flexibility is shown across framing conditions (left) and across repetitions (right) for both control (gray) and ASD participants (back).

Here again, there is no significant interaction between group and condition type (F[1,46]=0.55, p=0.46, R^2^=1.2%), but there is a significant main effect of condition type (F[1,46]=5.54, p=0.02, R^2^=10.7%) and a main effect of group (t[46]=3.4, p=0.001, R^2^=20.4). Post-hoc tests show that this group difference in repertoire’s flexibility is strong both across framings (t[46]=3.4, p=0.001, R^2^=20.7%) and across repetitions (t[46]=2.8, p=0.004, R^2^=14.4%). Also, ASD participants show no difference in repertoire’s flexibility when considered across framings or across conditions (p=0.26). This contrasts with control participants, who exhibit a significantly greater repertoire’s flexibility across framings than across repetitions (p=0.03).

If only, this computational analysis confirms that ASD participants are relatively insensitive to the game’s framing. But do these computational phenotypes provide clinically useful information, above and beyond performance scores? We address this question by assessing the accuracy of a diagnosis classifier relying upon peoples’ flexibility 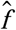 and ToM-sophistication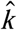. To begin with, we classified participants based upon computational phenotypes alone. The classifier only reached 67% of correct out-of-sample classifications. This is statistically significant (p=0.014), but worse than classification accuracy based upon performance patterns alone. However, when pooling performance patterns and computational phenotypes together, the classifier now yielded 79% of correct out-of-sample classifications (p<10^-4^). This is important, since it means that computational phenotypes bring additional, diagnosis-relevant, information.

Finally, we asked whether we could predict, from estimated computational phenotypes, inter-individual variations in symptom severity among ASD participants. More precisely, we focused on the ‘social’ and ‘stereotyped behavior’ subscores of the ADOS scale, which quantify social and non-social deficits, respectively. We found that inter-individual differences in computational phenotypes predict social deficits with high accuracy (F[4,15]=6.1, p=0.004, R^2^=62.1%) but not non-social deficits (F[4,15]=1.5, p=0.25, R^2^=28.8%).

## Discussion

In this work, we have performed a matched comparison of social and non-social behavioural adaptation in individuals with and without autism. In contrast to typically developed individuals, individuals with ASD do not change the way they play according to whether or not they believe they are competing with other humans. Typical individuals outperform people with ASD only when they think they are competing with another human being (and while playing against learning algorithms equipped with artificial mentalizing). However, people with ASD outperform typical individuals against non-mentalizing learning algorithms, irrespective of the task framing (social and non-social). The learning repertoire of individuals with ASD exhibits less flexibility and less ToM-sophistication, especially when the task is framed as a social game. Taken together, performance patterns and computational phenotypes correctly classify up to 79% of the participants according to their diagnosis. In addition, computational phenotypes predict 62% of the variance of the severity of social symptoms in ASD people.

Maybe the most striking result of our work is that people with ASD fail our “reverse Turing test”, in the sense that their cognitive strategy is insensitive to the game’s framing. Recall that we demonstrated this in four different ways: (i) ASD participants show no difference between performance or ToM-sophistication scores between framing conditions (cf. Fig. 2), (ii) performance variations induced by opponent types in different framing conditions are significantly correlated (see section 4 in Supplementary Text S1), (iii) model-free decompositions of their trial-by-trial choice sequences show no effect of framing (see section 5 in Supplementary Text S1), and (iv) their learning repertoire exhibits very low flexibility across framing conditions (cf. Fig. 4). Importantly, participants’ debriefing showed that the framing manipulation was similarly credible in both groups of subjects (see section 2 in Supplementary Text S1). In line with social motivational theories of autism [37], one may argue that, in contrast to control participants, ASD participants may not have been interested enough to invest the cognitive effort required for improving their performance in the social framing condition. Such global motivational and/or attentional interpretations are unlikely however, because ASD participants actually outperform controls against *0-ToM* in the social framing condition. In addition, financial incentive manipulations have no effect on performance in the game (see section 3 in Supplementary Text S1). In any case, our computational results rather suggest that people with ASD rely on a very limited learning repertoire, which they deem reliable in both social and non-social contexts. It is interesting to note that the model that best captures trial-by-trial choice sequences of ASD players, in both framing conditions, is the so-called “influence learning” strategy [33]. From a computational standpoint, this model possesses broad adaptive fitness because it essentially is a generic way of learning in reactive environments (i.e. environments that react to one’s actions). In other words, influence learning can be seen as an all-purpose cognitive toolkit that would be expected to perform well in a wide range of contexts, excluding challenging social interactions (cf. pattern of performances against RB, *0-ToM, 1-ToM* and *2-ToM* in section 9 in the Supplementary Text S1). Obviously, our experimental claim does not go as far as to assert that the cognitive repertoire of ASD people is generally limited to influence learning. Nevertheless, it provides a remarkable example of how people in the autism spectrum may solve the unavoidable trade-off between behavioural adaptability and cognitive complexity.

This type of trade-off is arguably steepest in ecological social contexts. Not only may subtle signals (e.g., facial expressions, speech prosody, etc…) reflect profoundly different mental states, but the stakes of typical social exchanges may be dynamic, partially implicit, multiple and even conflicting (e.g., impose a deal and induce sympathy). This implies that flexibility may be a critical feature of typical social cognition [38]. In this work, we provide two independent pieces of evidence in favour of this notion. First, in the ASD group, the severity of social symptoms is partially predicted by our measure of repertoire flexibility. Second, in the NT group, flexibility (between repetitions) is significantly higher in the social than in the non-social framing condition (see section 7 in Supplementary Text S1). On the one hand, these results contribute to the ongoing debate regarding the specificity of social cognition [23,39]. In brief, social cognition is special, if only because its flexibility is enhanced (notwithstanding its sophistication). On the other hand, they also bridge a gap between social and non-social theories of ASD. Recall that the latter typically take weak context-sensitivity as a feature of ASD cognition [40,41]. Importantly, our results tend to support the view that social deficits in ASD may be but a limiting case of the failure to account for the social context when drawing inferences about others.

## Methods

### Ethics statement

Behavioural assessments were performed in accordance with institutional ethical guidelines, which comply with the guidelines of the declaration of Helsinki. The research protocol was approved by the Ethical Committee of the Hôpital Rivière-des-Prairies, Montréal, where the tests were performed.

### Experimental methods

Participants: n=24 adults with ASD without mental nor language deficiency and n=24 NT control subjects participated in the study. All subjects were French speakers (Québec), and both groups were matched in terms of gender balance (ASD: 21 males, NT: 21 males), age (ASD: 25.5 y.o. ± 5.7; NT: 27.9 y.o. ± 8.6) and IQ (ASD: 104 ± 17; NT: 106 ± 14). ASD participants were assessed with ADOS-G and met DSM-5 criteria for ASD. NT participants went through a semi-structured interview to screen for any psychiatric treatment history, learning disorders, personal or family history (2 degrees) for mood disorder, ASD or schizophrenia. No included participant reported strong depressive symptoms (Beck depression Inventory score<20). All participants gave their informed consent, were fully debriefed at the end of the experiment, and received a financial compensation for their participation.

The behavioural task consists of a computerized game (60 trials each) with two framing conditions. In the *social* condition, the task was framed as an online competitive game with another participant. In the *non-social* condition, it was framed as a betting -casino-like-game. In fact, both games were played against four different learning algorithms with different artificial mentalizing sophistication (ranging from a random sequence with a bias to so-called *2-ToM* agents). At each trial, subjects had 1300 ms to make a binary choice (the place to hide or the slot machine to try), which was fed to the learning algorithms to compute online predictions of the participant’s action at the next trial. In total, each participant performed 2×4×2=16 games (2 framings, 4 opponent types, 2 repetitions) in a pseudo-randomized order. We refer the interested reader to the Supplementary Text S1 for more details regarding the experimental protocol.

### Computational modelling of learning strategies

In this section, we give a brief overview of the set of candidate learning models, with a particular emphasis on *k-ToM* models (because these are also used as on-line algorithms during the experimental phase). We will consider repeated dyadic (two-players) games, in which only two actions are available for each player (the participant and his opponent). Hereafter, the action of a given agent (resp., his opponent) is denoted by *a*^*self*^ (resp., *a*^*op*^). A game is defined in terms of its payoff table, whose entries are the player-specific utility *U (a*^*self*^, *a*^*op*^) of any combination of players’ actions at each trial. In particular, competitive social interactions simply reduce to anti-symmetric players’ payoff tables (see Table 1 below).

**Table 1:** Competitive payoff table. Participants play the role of the seeker, the opponent is the hider.

|  |  | Hider |  |
| --- | --- | --- | --- |
|  |  | Left | Right |
| Seeker | Left | 1,0 | 0,1 |
|  | Right | 0,1 | 1,0 |

By convention, actions *a*^*op*^ and *a*^*self*^ take binary values encoding the first (*a =*1) and the second (*a =* 0) available options. According to Bayesian decision theory, agents aim at maximising expected payoff *V = E [U* (*a*^*self*^, *a*^*op*^ )where the expectation is defined in relation to the agent’s uncertain predictions about his opponent’s next move. This implies that the form of the decision policy is the same for all agents, irrespective of their ToM sophistication. Here, we consider that choices may exhibit small deviations from the rational decision rule, i.e. we assume agents employ the so-called “softmax” probabilistic policy:

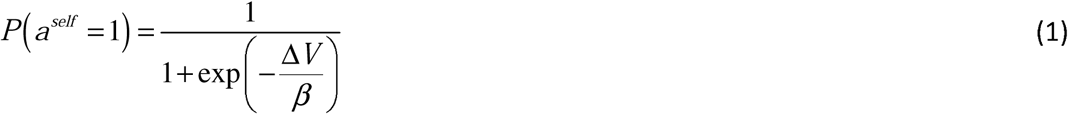

where *P* (*a*^*self*^ =1) is the probability that the agent chooses the action *a*^*self*^ = 1, Δ*V* is the expected payoff difference (between actions *a*^*self*^ =1 and *a*^*self*^ =0), and *β* is the so-called behavioural “temperature” (which controls the magnitude of deviations from rationality). The sigmoidal form of Equation 1 simply says that the probability of choosing the action *a*^*self*^ =1 increases with the expected payoff difference Δ*V*, which is given by:

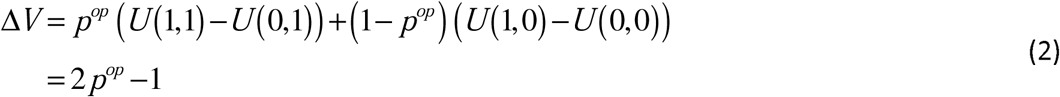

where *p*^*op*^ is the probability that the opponent will choose the action *a*^*op*^ =1, and the second line derives from inserting the above payoff matrix (Table1). In brief, Equation 2 simply says that participants are rewarded for correctly guessing where their opponent is hiding.

Let us now summarize the mathematical derivation of *k-ToM* models, which essentially differ in how they estimate *p*^*op*^ from the repeated observation of their opponent’s behaviour. We will see that *k* indexes a specific form of ToM sophistication, namely: the recursive depth of learners’ beliefs (as in “I believe that you believe that I believe…”). Note that k-ToM’s learning rule can be obtained recursively, starting with *0-ToM* [29].

By convention, a *0-ToM* agent does not attribute mental states to his opponent, but rather tracks his overt behavioural tendency without mentalizing. More precisely, *0-ToM* agents simply assume that their opponents choose the action *a*^*op*^ =1 with probability *p*^*op*^ =*s* (*x*_*t*_), where the unknown log-odds *x*_*t*_ varies across trials *t* with a certain volatility *σ* ^0^ (and *s* is the sigmoid function). Observing his opponent’s choices gives *0-ToM* information about the hidden state *x*, which can be updated trial after trial using Bayes rule, as follows:

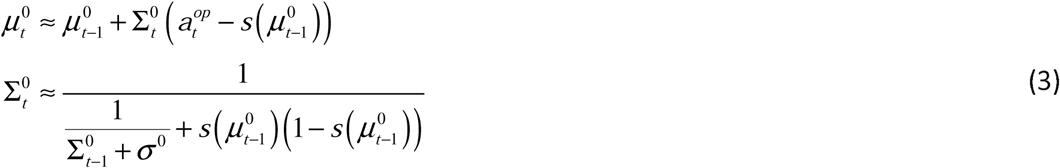

where 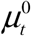 (resp. 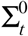) is the approximate mean (resp. variance) of *0-ToM*’s posterior distribution 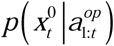. Inserting 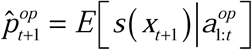into Equation 1 now yields *0-ToM*’s decision rule. Here, the effective learning rate is the subjective uncertainty ∑^0^, which is controlled by the volatility *σ* ^0^. At the limit *σ* ^0^ ⟶ 0, Equation 3 converges towards the (stationary) opponent’s choice frequency and 0-ToM essentially reproduce “fictitious play” strategies [34].

*0-ToM*’s learning rule is the starting point for a 1-ToM agent, who considers that she is facing a *0-ToM* agent. This means that *1-ToM* has to predict *0-ToM*’s next move, given his beliefs and the choices’ payoffs. The issue here is that *0-ToM*’s parameters (volatility *σ*^0^ and exploration temperature *β*) are unknown to *1-ToM* and have to be learned, through their non-trivial effect on *0-ToM*’s choices. At trial *t* +1, a 1-ToM agent predicts that *0-ToM* will chose the action *a*^*op*^ =1 with probability 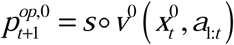, where the hidden states 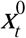 lumps *σ*^0^ and *β* together and the mapping *v*^0^ is derived from inserting *0-ToM*’s learning rule (Equation 3) into Equations 1-2. Similarly to *0-ToM* agents, 1-ToM assumes that the hidden states 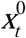vary across trials with a certain volatility *σ*^1^, which yields a meta-Bayesian learning rule similar in form to *0-ToM*’s, but relying on first-order meta-beliefs (i.e. beliefs about beliefs). In brief, 1-ToM eventually learns how her (0-ToM) opponent learns about herself, and acts accordingly (cf. Equations 1-2).

*1-ToM* agents are well equipped to deal with situations of observational learning. However, when it comes to reciprocal social interactions, one may benefit from considering that others are also using ToM. This calls for learning strategies that rely upon higher-order meta-beliefs. By construction, *k-ToM* agents (*k ≥* 2) consider that their opponent is a *κ* -*ToM* agent with a lower ToM sophistication level (i.e.: *κ* < *k*). Importantly, the sophistication level *κ* of *k-ToM*’s opponent has to be learned, in addition to the hidden states *x*^κ^ that control the opponent’s learning and decision making. The difficulty for a k-ToM agent is that she needs to consider different scenarios: each of her opponent’s possible sophistication level *κ* yields a specific probability 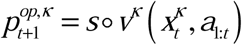 that she will choose action *a*^*op*^ =1 The ensuing meta-Bayesian learning rule entails updating *k-ToM*’s uncertain belief about her opponent’s sophistication level *κ* and hidden states *x*^*κ*^:

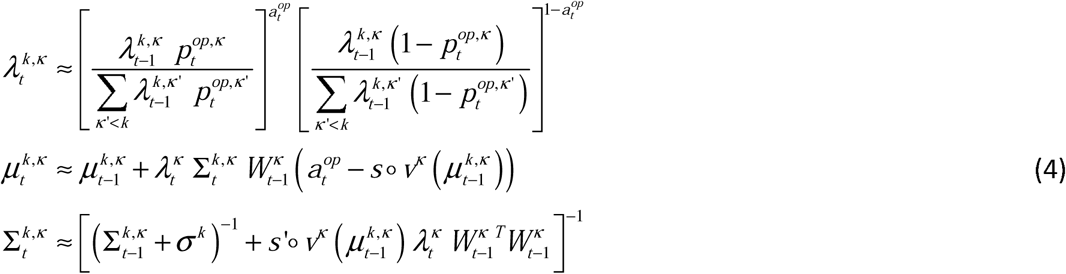

where 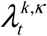 is *k-ToM*’s posterior probability that her opponent is *κ*-ToM, and *W* ^*κ*^ is the gradient of *v*^*κ*^ with respect to the hidden states *x* ^*κ*^. Equation 4 also captures *1-ToM*’s learning rule, when setting 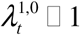⁏ 1. Note that although the dimensionality of k-ToM’s beliefs increases with *k, k*-*ToM* models do not differ in terms of the number of their free parameters. More precisely, *k-ToM*’s learning and decision rules are entirely specified by their prior volatility *σ* ^*k*^ and behavioural temperature *β*.

Formally speaking, only *k-ToM* agents with *k*≥1 are mentalizing about others’ covert mental states, i.e. represent and update others’ beliefs. They can do this because they adopt the intentional stance [32], i.e. they assume that *p*^*op*^ is driven by their opponent’s hidden beliefs and desires. More precisely, they consider that the opponent is himself a Bayesian agent, whose decision policy *p* ^*op*^ =*P*(*a* ^*op*^ =1) is formally similar to Equation 1. This makes *k*-ToM meta-Bayesian learners [35] that relies upon recursive belief updating (“I believe that you believe that I believe…”). Critically, the recursion depth k induces distinct ToM sophistication levels, whose differ in terms of how they react to the history of players’ actions in the game.

With the exception of *0-ToM*, we so far only described sophisticated learning models that are capable of (artificial) ToM. But clearly *0-ToM* is not the only way people may learn in social contexts without mentalizing. We thus consider below other learning strategies that may populate peoples’ learning repertoire.

First, let us consider a heuristic learning model, whose sophistication somehow lies in between *0-ToM* and *1-ToM*. In brief, “influence learning” adjusts a *0-ToM*-like learning rule to account for how her own actions may influence her opponent’s behaviour [33]:

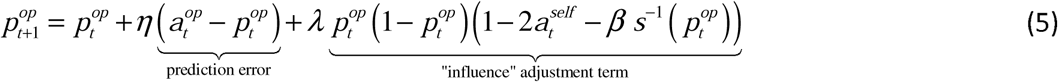

where η (resp. *λ*) controls the relative weight of its prediction error (resp. the “influence” adjustment term). Numerical simulations show that, in a competitive game setting, Inf wins over *0-ToM* but loses against *k-ToM* players with k≥1. In other terms, although it is in principle able to adapt to reactive environments, Inf cannot successfully compete with learners endowed with mentalizing [28].

Second, participants may learn by trial and error, eventually reinforcing the actions that led to a reward. Such learning strategy is the essence of classical conditioning, which is typically modelled using reinforcement learning or RL [42]. In this perspective, participants would directly learn the value of alternative actions, which bypasses Equation 2. More precisely, an RL agent would update the value of the chosen option in proportion to the reward prediction error, as follows:

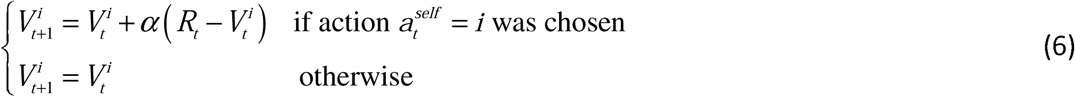

where 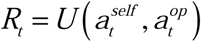is the last reward outcome and *α* is the (unknown) learning rate. At the time of choice, RL agents simply tend to pick the most valuable option (cf. Equation 1).

Third, an even simpler way of adapting one’s behaviour in operant contexts such as this one is to repeat one’s last choice if it was successful and alternate otherwise. This can be modeled by the following update in action values:

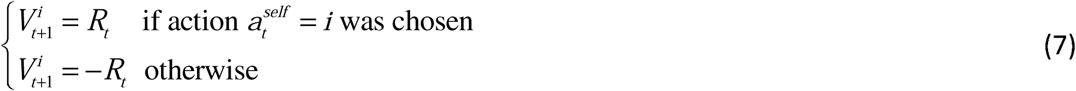

This strategy is called win-stay/lose-switch (WS), and is almost identical to the above RL model when the learning rate is *α* = 1. Despite its simplicity, WS can be shown to have remarkable adaptive properties [43].

Last, the agent may simply act randomly, which can be modeled by fixing the value difference to zero (Δ*V =* 0). Although embarrassingly simple, this probabilistic policy eventually prevents one’s opponent from controlling one’s expected earnings. It thus minimizes the risk of being exploited at the cost of providing chance-level expected earnings. It is the so-called “Nash equilibrium” of our “hide and seek” game. Since we augment this model with a potential bias for one of the two alternative options (as all the above learning models), we refer to it as *biased Nash or BN*.

### Empirical estimates of computational phenotypes

Our working hypothesis is that people may not always rely on the same learning model across different game sessions or conditions. Rather, they select a learning strategy from among a repertoire, whose flexibility and ToM sophistication define our computational phenotypes. The empirical estimation of these thus consists of three steps. First, we perform a statistical (Bayesian) comparison of learning models [44]. For each subject, we fit trial-by-trial actions sequences *a*_1:60_ with each learning model (*m* ∈{*BN, WSLS, RL, 0-ToM, Inf, 1-ToM, 2-ToM, 3-ToM})* using a variational-Laplace approach [45,46]. This eventually yields 8×48×4×2×2=6144 model evidences *p*(*a*_1:60_❘*m)* (8 models, 48 participants, 4 opponent conditions, 2 framing conditions, 2 repetitions).

Second, we define the *repertoire*’s *flexibility* 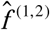(between conditions 1 and 2) in terms of the posterior probability that a given participant employs different learning strategies across two conditions:

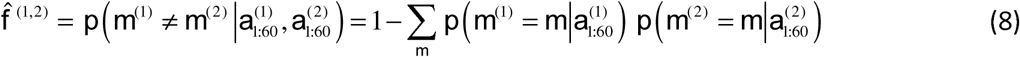

where *m*^(1)^ (resp. *m*^(1)^) is the participants’ learning strategy in the first (resp. second) condition, 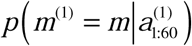(resp. 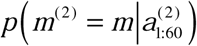) is the probability that the participant had a learning strategy *m* given his trial-by-trial choice sequence 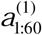(resp. 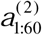in condition 1 (resp. condition 2). Note that we measure the repertoire’s flexibility 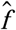both across framings and across repetitions.

Third, we define the *repertoire*’s *ToM-sophistication* 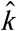in terms of the expected depth of recursive belief update:

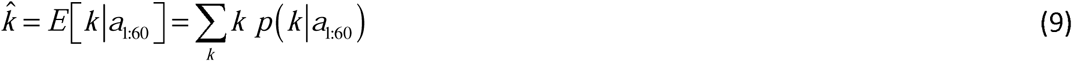

where *p*(*k❘a*_1:60_) = *p*(*m* = “*k − ToM*“❘*a*_1:60_) is the posterior probability of model *k-ToM* given the participant’s trial-by-trial choice sequence *a*_1:60_. Note that we restrict the summation in Equation 9 to *k-ToM* models, because the depth k of recursive beliefs is not defined for the other learning models. Note that we measure the *repertoire*’s *ToM-sophistication* 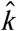in both framing conditions (social and non-social).

All statistical analyses were performed using the VBA toolbox [36].

## Acknowledgements

Authors thank Alexandra Duquette and Patricia Jelenic for contributing with data collection, and Pr. Laurent Mottron for enabling the recruitment of ASD patients.

## Financial disclosure

BFA acknowledges support from “Fondation Les Petits Trésors de l’Hôpital Rivière des Prairies” and Fonds de Recherche en Santé du Québec (FRQS). MD and JD have nothing to declare.

